# Focused learning by antibody language models using preferential masking of non-templated regions

**DOI:** 10.1101/2024.10.23.619908

**Authors:** Karenna Ng, Bryan Briney

## Abstract

Existing antibody language models (**LMs**) are pre-trained using a masked language modeling (**MLM**) objective with uniform masking probabilities. While these models excel at predicting germline residues, they often struggle with mutated and non-templated residues, which are crucial for antigen-binding specificity and concentrate in the complementarity-determining regions (**CDRs**). Here, we demonstrate that preferential masking of the non-templated CDR3 is a compute-efficient strategy to enhance model performance. We pre-trained two antibody LMs (**AbLMs**) using either uniform or preferential masking and observed that the latter improves residue prediction accuracy in the highly variable CDR3. Preferential masking also improves antibody classification by native chain pairing and binding specificity, suggesting improved CDR3 understanding and indicating that non-random, learnable patterns help govern antibody chain pairing. We further show that specificity classification is largely informed by residues in the CDRs, demonstrating that AbLMs learn meaningful patterns that align with immunological understanding.

## INTRODUCTION

Antibodies are a crucial component of the humoral immune system. Their massive diversity, estimated to be as high as 10^18^ unique antibodies,^1^ confers the potential to bind and neutralize any non-self antigen with remarkable specificity. The pre-immune antibody repertoire has combinatorial diversity from independent V(D)J recombination of both the heavy and light chains in each B-cell, followed by pairing of those chains as a dimer of heterodimers. Each chain has three complementarity-determining regions (**CDRs**) which form loops that largely determine antigen-binding specificity. Upon infection, antigen-specific antibodies undergo further affinity maturation, where mutations are stochastically introduced and selected for their ability to strengthen binding affinity. Although somatic mutations can be found throughout the antibody gene, they are concentrated in the CDRs.

The amino acid sequence of a protein encodes its structure and function, analogous to how the order and context of the words in a sentence encode its meaning. This parallel has inspired the adaptation of transformer-based language models (**LMs**),^2^ originally developed for natural language processing (**NLP**), for analysis of protein sequences. Protein LMs (**PLMs**) have shown success at understanding evolutionary fitness,^3^ as well as protein structure and function.^4,5^ Antibody LMs (**AbLMs**), pre-trained primarily or exclusively on antibody sequence data, have outperformed general PLMs at learning antibody-specific features such as affinity maturation,^6^ antigen specificity,^7^ and paratope position.^8^ However, both PLMs and AbLMs struggle to learn patterns beyond what is germline-encoded. This is most evident in the poor performance in prediction of the CDR3,^7,9^ which spans the junction between germline V(D)J gene segments and is enriched with non-templated mutations.

Many PLMs and AbLMs use a BERT-like architecture to generate sequence embeddings that can be applied to downstream tasks, such as structure prediction^5,10^ or directed evolution to improve binding affinity.^3^ These models are trained using a masked language modeling (**MLM**) objective, where a set of “masked” tokens are predicted based on the remaining bidirectional context.^11^ Prediction of these masked tokens, which we call “training signal,” is then used to iteratively optimize the model weights. BERT randomly masks 15% of the tokens in each input sequence, and this uniform masking rate has been adopted by most BERT successors, regardless of model size or specific architecture.^4–9,12^ However, this may not be optimal for training AbLMs.

Firstly, the 15% masking rate is thought to balance sequence corruption with training efficiency: overmasking of tokens results in limited context and poor representation learning, but pre-training is inefficient when too few tokens are predicted. However, higher masking rates have been shown to improve LM performance on multiple NLP benchmarks, especially for BERT-large sized models.^13^ Increasing the masking rate results in the model predicting more tokens for each training example, effectively learning from more of the sequence. This efficient use of training data is especially relevant for AbLMs, which are limited by a lack of antibody sequence training data at scale.^7,9,14^ Secondly, both pre-immune and affinity-matured antibody sequences consist predominantly of germline-encoded (templated) residues. Thus, under uniform masking, the majority of masked residues are in templated regions.^9^ This “frequency-bias problem” limits pre-training efficiency, lowers the quality of the representations of these rare and valuable tokens,^15,16^ and makes it difficult for the model to learn non-templated patterns.

Alternative masking strategies have been explored for NLP, including changing overall masking rates^13^ and frequency-weighted token sampling.^15^ To our knowledge, alternative masking has not been applied to AbLM training, which is surprising since the modular nature of antibody recombination creates relatively distinct regions of high complexity (non-templated) and low complexity (templated or germline-encoded). The goal of this study was to determine to what extent AbLMs can improve when trained using an alternative masking strategy developed specifically for antibody sequence data.

Here, we present pre-training with preferential masking of the non-templated CDR3. We hypothesized this strategy to be more informative for immunological downstream tasks, since training signal is increased from the antibody region with the most diversity (difficult to learn) and most relevance to antigen-binding. To explore this, we pre-trained two AbLMs with either uniform or preferential masking. We demonstrate that the preferential model shows enhanced ability to distinguish native chain pairings from random pairings and identify binding specificity, both of which are immunologically relevant features of antibodies. Using explainable artificial intelligence (**XAI**), we further show that classification by binding specificity is largely informed by residues in the CDRs, suggesting that our models are learning meaningful patterns that align with immunological understanding of antibody functional regions.

## RESULTS

### Preferential masking implementation

Most models trained with an MLM objective follow the original BERT implementation, where 15% of tokens are independently and randomly selected for masking under a uniform probability distribution.^11^ Since model weights are only optimized based on masked token predictions, it follows that model performance would be sensitive to the specific tokens that were masked. Our preferential masking strategy increases the fraction of masked tokens that come from the CDR3, a diverse, non-templated region which has been shown to be difficult for models to learn.^7,9^ This is achieved by increasing the masking probability to 25% in the CDR3s of both the heavy and light chains. To enable comparison against the conventional uniform masking strategy and rule out increased overall masking as a confounding factor, the average masking probability across the entire input sequence length is maintained to be 15%. As a result, tokens outside of the CDR3s are masked at a rate of <15%. An overview of our preferential masking strategy is shown in ***Fig. 1***, and implementation details can be found in the methods.

**Figure 1.**
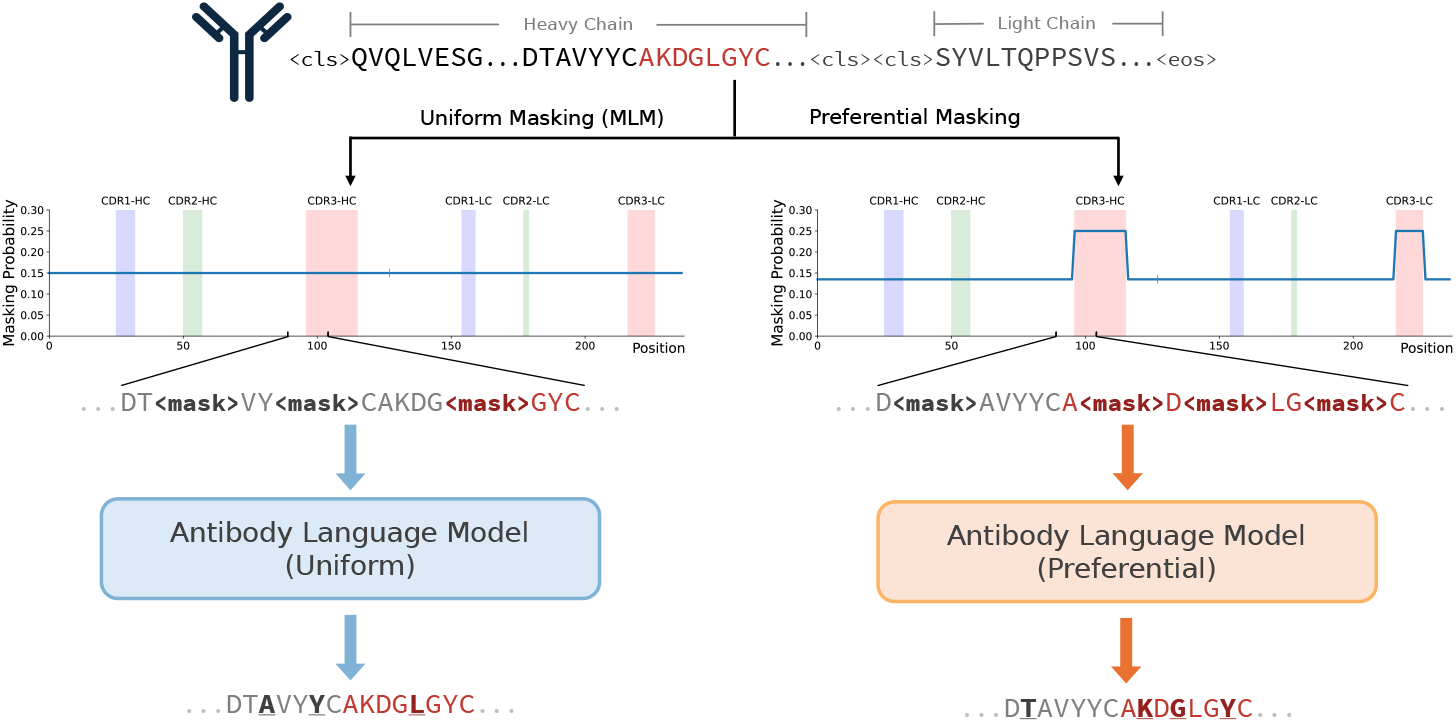
Uniform versus preferential MLM methodology. Conventional masking strategies (left) employ a uniform masking probability of 15% over the entire input sequence. Preferential masking (right) increases the masking probability in the CDR3s (shown in red) to 25% while maintaining the 15% average masking probability over the entire input sequence length.

### Preferential masking enables more efficient model pre-training

We pre-trained two masked AbLMs using Dataset A – one using conventional uniform masking, and one using our preferential masking strategy. Pre-training on paired sequences was chosen to enhance learning of cross-chain features.^7^ Both used the same ESM-2-based model architecture, which includes rotary position embeddings and pre-layer normalization.^17,18^ An encoder-only architecture was chosen to allow for the generation of sequence embeddings that can be used for downstream tasks such as sequence classification.

The validation loss of the preferential masking model converged with 40% less training time than the uniform model (***Fig. 2A***), suggesting that preferential masking enables learning of the same amount of information with fewer passes through the data. To identify if improvements in the CDRs were driving this change, validation loss was separated by antibody region (non-CDR, CDR1, CDR2, and CDR3). In all regions, the minimum loss of the preferential model was comparable or less than that of the uniform model (***Fig. 2B-E***), with the CDR3 loss additionally converging earlier (***Fig. 2E***). This suggests that preferential masking allows the model to extract more information from the sequence, and that reduced masking frequency outside of the CDR3 does not negatively impact representation learning. Overall validation loss of the uniform and preferential models began to increase at 350,000 and 250,000 steps, respectively (***Fig. 2A***), while the training loss continued to decrease (not shown), indicating overfitting and informing model checkpoint selection.

**Figure 2.**
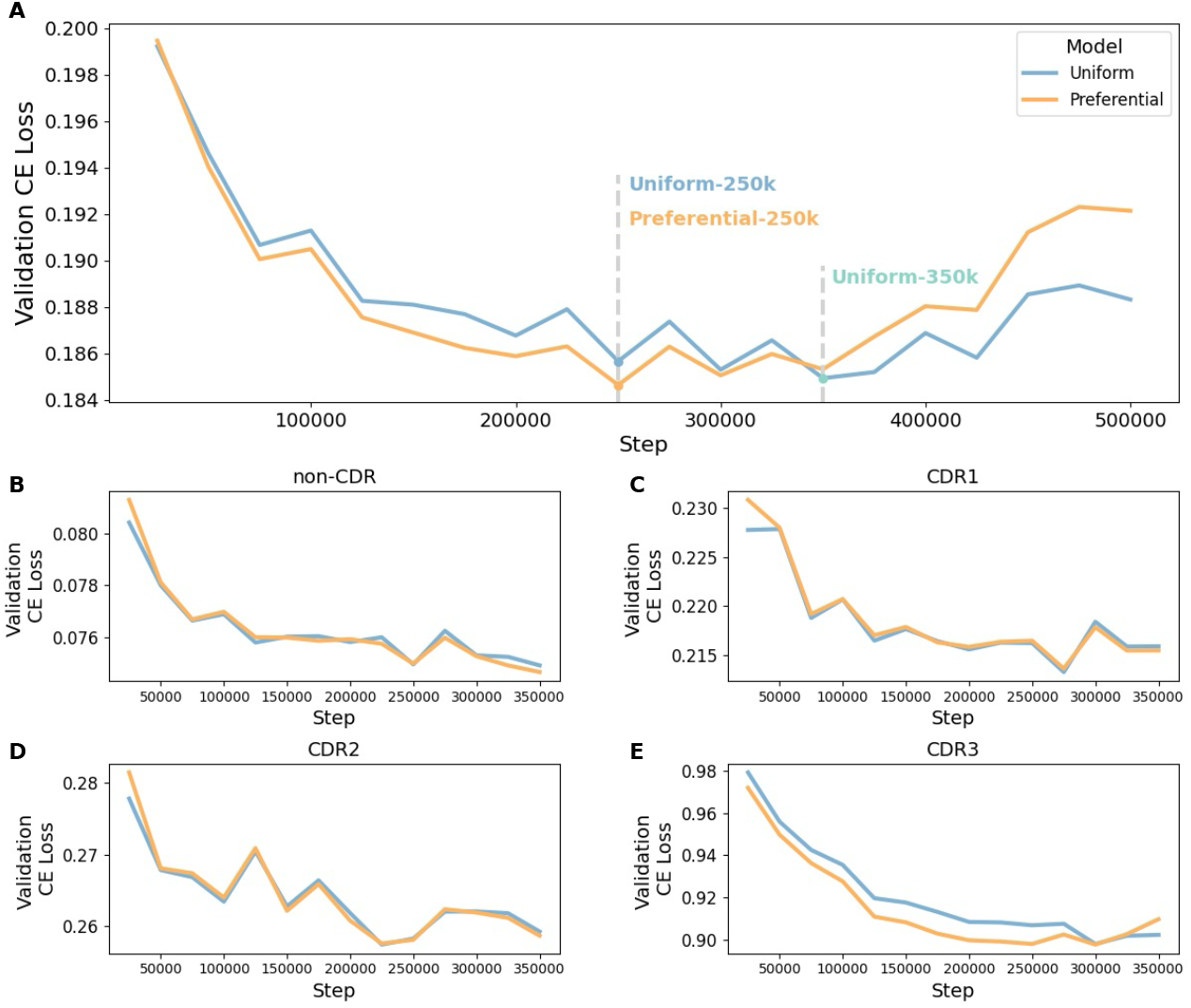
Pre-training validation loss curves. (***A***) Base model validation loss over the course of pre-training. Model checkpoints chosen for downstream analyses are indicated by the dashed lines. Overall validation loss is separated into non-CDR (***B***), CDR1 (***C***), CDR2 (***D***), and CDR3 (***E***) to examine progress of each region. The optimal checkpoints at 250,000 and 350,000 steps were used for downstream analyses, so regional loss plots are limited to the relevant training steps.

In the subsequent analyses, we evaluate the uniform and preferential masking models through two comparisons. First, we assess the models at their lowest error rate on the validation dataset: the uniform model at 350,000 steps (Uniform-350k), and the preferential model at 250,000 steps (Preferential-250k). Second, we compare the models trained with the same computational resources: Preferential-250k versus Uniform-250k.

### Preferential masking improves representation learning of mutated sequences

Antibody sequence diversity is highest in the CDR3, which contains the junction between germline V, D, and J segments, and non-templated sequence addition and deletion produces variable lengths and increased complexity.^9,10^ Preferential masking amplifies training signals from the CDR3, so we expect better learning of non-templated patterns when compared to the conventional uniform strategy.

Step-matched model performance was assessed by separately analyzing the per-position prediction accuracy of 1000 unmutated and mutated sequences from Dataset B. Grouping predictions by their corresponding framework region (**FR**) or CDR revealed much stronger model performance on unmutated sequences (***Fig. 3A***), which was expected since these are germline-encoded and contain less complexity than their mutated counterparts. In both sequence types, the CDR3s were observed to have considerably weaker performance than the FRs, consistent with previous analyses that germline-encoded features are more readily learned by LMs than complex patterns found in non-templated, junctional regions.^7,9^ After Bonferroni correction for multiple testing (14 regions), the Preferential-250k model performed significantly better than the Uniform-250k model at predicting residues in the mutated CDRH3 (p = 2.53×10^−9^) and CDRL3 (p = 3.52×10^−3^) (***Fig. 3B***), suggesting that preferential masking is recovering predictions of somatic mutations (introduced during affinity maturation). Though residues in the conserved framework regions were masked at a lower rate, we observed no significant decrease in performance, indicating that germline-encoded regions can be effectively learned with less training signal. The uniform model with 40% more training (Uniform-350k) reaches the Preferential-250k model’s performance in the CDR3 and performs slightly better in some non-CDR3 regions (***Fig. S1***).

**Figure 3.**
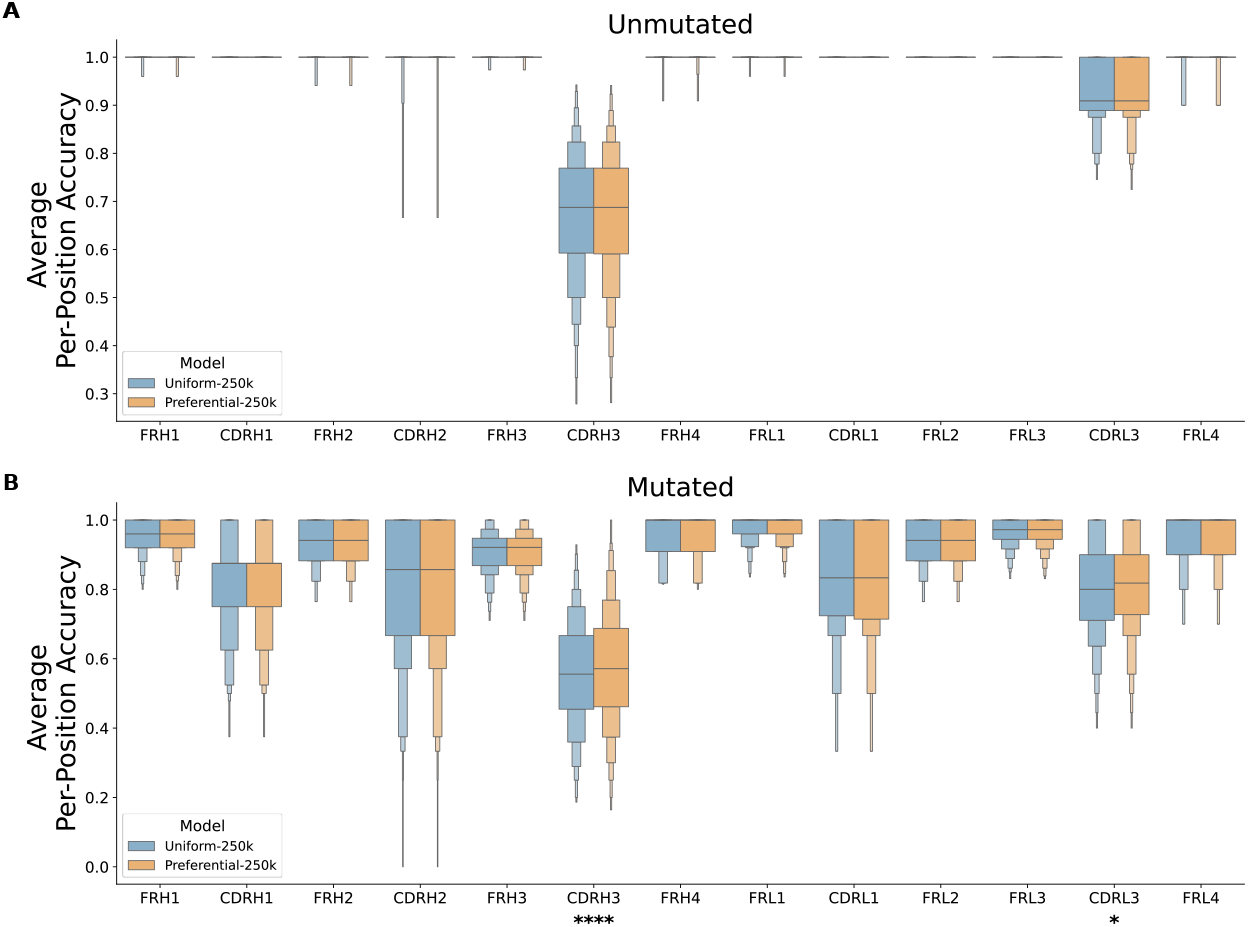
Per-position residue prediction accuracy. For 1000 unmutated (***A***) and mutated (***B***) test sequences, each residue was iteratively masked and predicted by both models (Uniform-250k and Preferential-250k). Mean prediction accuracy is plotted for each FR and CDR. Statistical significance for each region was calculated using a two-sided paired t-test with Bonferroni correction for multiple testing (14 regions).

### Productive VH/VL pairing is informed by learnable, non-random patterns

Antibody heavy and light chains independently undergo V(D)J recombination before their variable domains, **VH** and **VL**, respectively, pair to form a dimer. The interaction between the VH and VL is important for determining antibody stability and has been found to affect peptide binding kinetics,^19^ as the antigen-binding region is composed of CDR loops on both chains.^10^ VH/VL pairing is thought to be largely random, and suggested pairing preferences are not well understood.^19,20^ Given the crucial interaction between the VH and VL in the CDRs, preferential masking of the CDR3s may lead to improved learning of features that determine productive VH/VL pairs.

To assess this, we trained a sequence classification head to identify native vs. shuffled VH/VL pairs. We performed this test on two class-balanced datasets: Dataset C (∼65,000 sequence pairs), and Dataset D (∼140,000 sequence pairs).

All models classified native vs. shuffled pairs with over 60% accuracy, indicating at least some factors governing VH/VL chain pairing are non-random. The Preferential-250k model outperformed uniform models across metrics (Accuracy, AUC, AUPR, MCC) (***Fig. 4A***) and maintained superior performance throughout training (***Fig. 4B-C***), suggesting its embeddings were more informative for specificity classification. Most absolute metrics were higher for the classifier models trained on Dataset C, possibly due to inherent similarities to the pre-training data (Dataset A) as described in the methods. In addition, this dataset included both naïve and memory B-cell sequences. AbLMs have been shown to distinguish between unmutated (naïve) and mutated (memory) sequences.^7,21^ To examine if this distinction may have confounded pair classification, Dataset C predictions were separated into two groups: pairs where both chains were unmutated or mutated (“Same”), and pairs where only one of the two chains was mutated (“Different”). We observed that prediction accuracy was much higher for the “Different” pairs (***Fig. 4D***), indicating that shuffled pair predictions were likely skewed because the models had identified differentiating features between unmutated and mutated sequences. Dataset D, composed of only memory B-cell sequences from distinct donors from Dataset A, was designed to remove these confounding factors, and the increased difficulty of this task is reflected by the lower performance metrics. However, the preferential masking model’s improved performance still holds, suggesting that productive VH/VL pairing is informed by patterns in the CDR3 and that increased pre-training signal from this region improved learning of these patterns.

**Figure 4.**
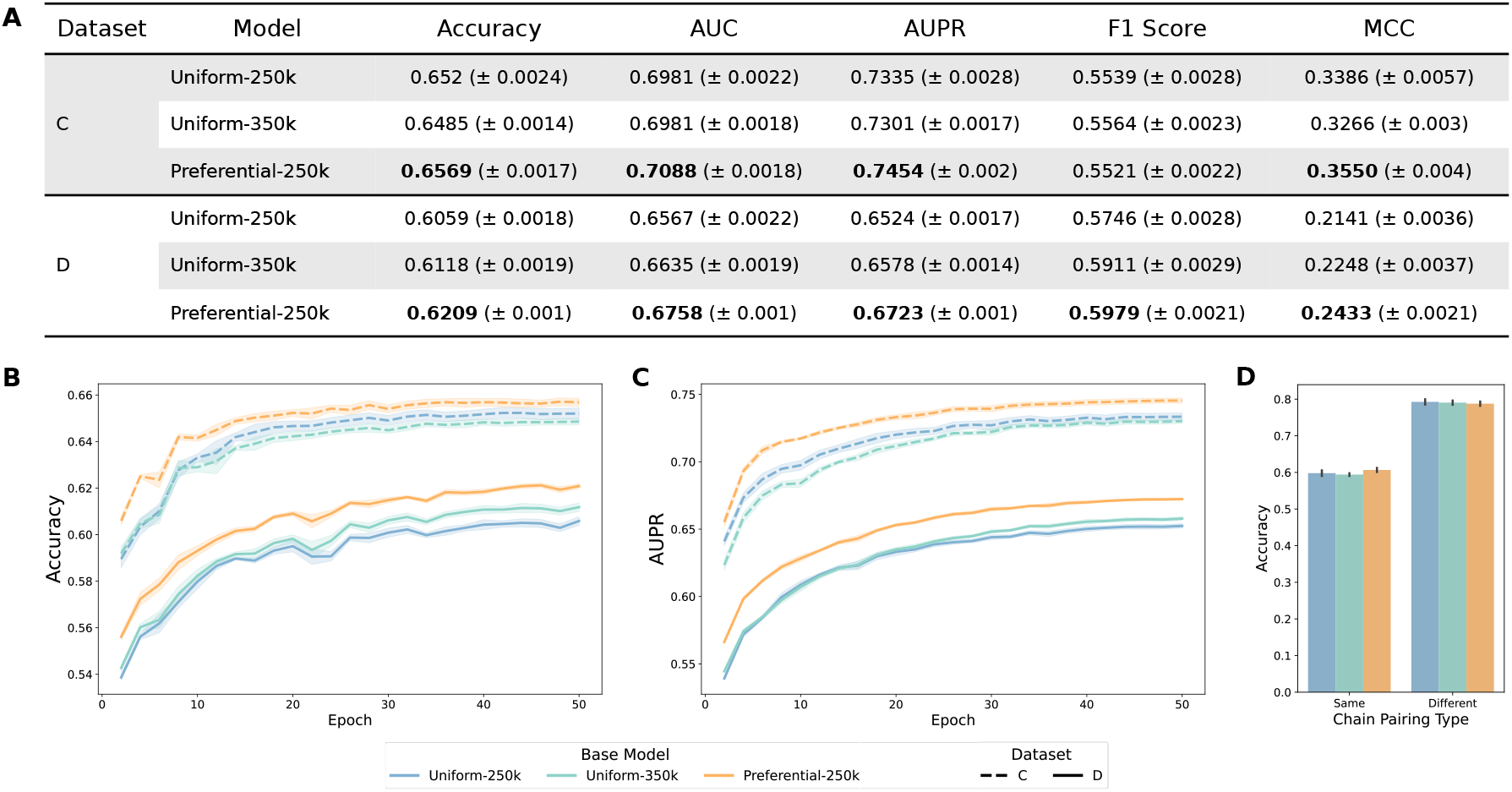
Native vs. shuffled chain pairing classification. (***A***) Test set metrics for pair classifier models at the end of training. Accuracy (***B***) and AUPR (***C***) over the course of classifier head training. (***D***) Accuracy for Dataset C separated by chain pairing type: the heavy and light chains are either both unmutated or mutated (“Same”), or only one of the chains is mutated, while the other is unmutated (“Different”). Mean and SE is shown for 5 independent training runs using 5-fold CV, and the highest values that are statistically significant for each dataset are bolded.

### Pre-training with preferential masking improves binding specificity classification

Previous studies have shown that AbLMs are able to classify sequences by antigen-binding specificity.^7,14^ Though the antibody repertoire is very diverse, antibodies frequently utilize recurring sequence motifs to target a specific epitope, even across individuals.^14^ These motifs are often located in the CDR3 because of its longer loop structure, high variability, and accumulation of mutations during affinity maturation.^9,10^ Given the critical role of the CDR3 in the antibody-antigen interaction, preferential masking of this region may enhance the model’s ability to classify sequences by antigen-binding specificity.

To explore this, we trained a specificity classification head on Dataset E (∼25,000 paired sequences) for binary classification of antibodies as CoV-specific or not. The negative set (non CoV-specific) was composed of randomly selected memory B-cell sequences obtained pre-2020 from several healthy donors to minimize false negative classifications.

We observed that the Preferential-250k model outperformed both uniform models on most metrics (***Fig. 5A***), highlighting the importance of the CDR3 in determining binding specificity. This improvement was consistent over the course of training (***Fig. 5B-C***), suggesting that the output embeddings by the preferential masking base model were more informative for specificity classification than those from the uniform model.

**Figure 5.**
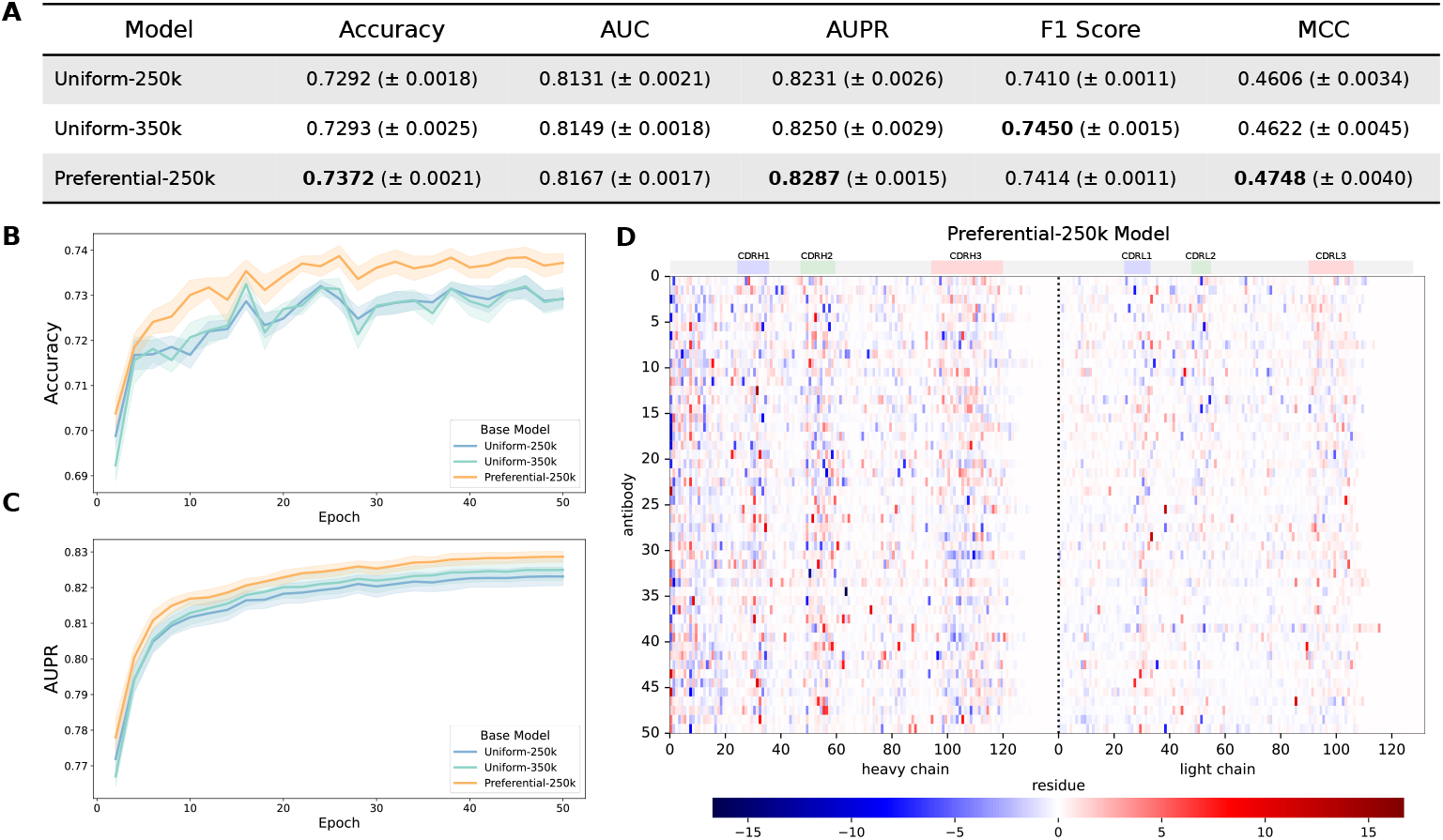
CoV-specificity binary classification and AttCAT analysis. (***A***) Test set metrics for CoV classifier models at the end of training. Accuracy (***B***) and AUPR (***C***) over the course of classifier head training. Mean and SE is shown for 5 independent training runs using 5-fold CV, and the highest values that are statistically significant for each dataset are bolded. (***D***) Normalized AttCAT impact scores with respect to the correct label class for 50 systematically chosen test sequences for the CoV specificity classifier trained on the Preferential-250k base model. Top color bars indicate approximate CDR locations. Sequences are sorted in ascending order by average prediction probability across the classifiers trained on all 3 base models.

### Models identify residues in the CDRs as important for determining binding specificity

One major limitation of LLMs in both the NLP and protein spaces is interpretability.^22^ As models grow in complexity and are increasingly used to make healthcare decisions, it is important to develop XAI methods to enable better human understanding of model reasoning. We sought to understand the classifier models’ decision-making process for identification of CoV-specific sequences. Since the antigen-binding region is largely composed of the CDR loops, we expect that residues in the CDRs would be identified as most salient for determining binding specificity.

A common method of explaining a transformer’s output is to plot the relative importance distribution over the input tokens using raw attention weights from single or multiple layers. While this technique has been applied to existing AbLMs,^7,8^ it does not consider information flow via other paths, such as through the residual connections, and does not consistently correlate with model performance or other feature importance indicators.^23^ Thus, we used the gradient-based XAI technique Attentive Class Activation Tokens (**AttCAT**)^23^ to explore which residues were model-identified as “important” for classifying a sequence as CoV-specific.

AttCAT impact scores were computed for 50 systematically chosen test sequences from Dataset E for each classifier model with respect to the correct label class (***Fig. 5D, Fig. S2***). Positive scores (shown in red) indicate importance towards the CoV-positive class, while negative scores (shown in blue) indicate importance towards the CoV-negative class. High importance residues, as indicated by score magnitude (darker colors), were concentrated in the CDRs with stronger activation in the heavy chain, suggesting that the models have learned that these are the key regions for determining binding specificity. This is consistent with previous XAI analyses on PLMs by Wenzel et al.,^24^ which showed that sequence regions model-identified as “important” were correlated with known functional regional annotations.

## DISCUSSION

Advancements in NLP have been applied to PLMs and AbLMs with little adaptation of the methods to accommodate the unique features of biological data. While this approach has enhanced our understanding of general structure and function, these models fall short in capturing the highly variable CDR3, which contains the junction between V(D)J segments and plays a crucial role in antigen-binding. Here, we sought to improve CDR3 understanding by implementing an antibody-specific modification to the pre-training strategy. While prior studies have explored modifications to the training data,^7,14^ few^9^ have approached this problem from the pre-training strategy.

We present preferential masking as a compute-efficient method for antibody representation learning, achieving predictive performance similar to uniform masking with 40% less training time (***Fig. 2, 3***). Our use of dynamic masking^12^ effectively presents a unique sequence context for each input during each epoch, demonstrating that preferential masking also enhances data efficiency, which is particularly important given the limited scale of training data available for AbLMs.^7,9,14^ Increasing the frequency of masking and prediction in high-diversity regions also increases the diversity of the training data observed by the model, leading to greater efficiency. Our results indicate that preferential masking enhances downstream classification of antibody sequences by native chain pairing (***Fig. 4***) and binding specificity (***Fig. 5***), highlighting the role of the CDR3 in antibody development and function. While these specific binary classification tasks may have limited practical use, our findings indicate that preferential masking enhances the learning of immunologically relevant patterns, potentially enhancing performance on other downstream tasks such as structure prediction.

Analysis of native vs. shuffled pair classification indicates that antibody VH/VL pairing is not random, contrasting previous studies^20^ and supporting emerging evidence of potentially learnable patterns that influence pairing decisions.^25^ This observation is particularly notable for Dataset D, which contains only memory B-cell sequences from donors distinct from those used in pre-training. This suggests that non-random pairing is a true biological phenomenon rather than a result of confounding factors. Insights from these models may improve strategies for expressing more stable antibodies for use in laboratory settings.

AttCAT analysis reveals that our CoV-specificity classifier models are primarily informed by residues in the CDRs, especially on the heavy chain (***Fig. 5D***). The antigen-binding pocket consists of CDR loops from both heavy and light chains, with the CDRH3 being the most diverse and influential in driving specificity.^10^ This indicates that our models have correctly identified the crucial regions for determining binding specificity, validating our use of LM techniques for antibody sequence modeling. These results suggest that preferential masking truly does enable better learning of immunological features.

Entropy or importance-based weighting has been successfully applied to NLP tasks, leading to improved training efficiency and downstream performance.^15^ The non-templated CDR3 represents areas of high entropy, selected for preferential masking due to its functional relevance and significantly higher diversity. Though our approach is limited by the need for data to be annotated with the locations of non-templated regions, many antibody sequencing pipelines such as CellRanger^26^ and abstar^27^ already include these annotations. Also, minimal additional computation is required to derive the masking probabilities, as this can be done with only a single pass through the dataset. Thus, integrating preferential masking into existing AbLM training is relatively straightforward. However, a more sophisticated weighted masking strategy could further enhance efficiency and reveal additional significant areas for improving antibody representation learning.

Although non-templated residues are primarily found in the CDR3, they are not exclusive to this region; other existing sequence annotations, such as mutations resulting from affinity maturation, could be the targets of preferential masking. Further, a dynamic, data-driven strategy could make this process entirely self-supervised. For example, focal loss, which adjusts token impact on the loss by its prediction confidence at each pre-training step, has been explored for AbLMs and shown to increase out-of-distribution predictions.^9^ A similar approach could be adapted for weighting the masking probability.

Our classifier model analyses indicate that there are learnable, sequence-inherent patterns that can effectively differentiate groups of antibodies. These are likely concentrated in the CDR3, as evidenced by the improved performance of classifiers trained on the preferential base model. A better understanding of these patterns could deepen our knowledge of the fundamental immunological properties governing antibody development and function. However, deciphering the decision-making processes of LMs remains challenging. This study is among the first^14^ to apply XAI techniques to antibodies, illustrating how LM-learned patterns can become interpretable. While AttCAT offers a high-level overview of information flows and learned parameters, it lacks the granularity needed to extract detailed patterns. Additionally, because AttCAT is designed for classification models, its insights are limited to task-specific contexts and do not fully capture the knowledge acquired by the underlying pre-trained models. Further work on XAI tailored to biological data is needed to enable discovery of novel patterns from complex LMs.

There is still significant potential for improving AbLMs. These models are currently trained on relatively small datasets compared to state-of-the-art LMs for NLP. Expanding the training data via additional sequencing or mining of sequence databases like the Observed Antibody Space (**OAS**)^28^ could enhance the diversity of learned patterns and improve generalizability. Furthermore, incorporating additional modalities may enhance model performance. For example, models like ESM3, which combine sequence data with structural and functional information, have demonstrated improvements in generative capabilities.^29^ Given that AbLMs outperform general PLMs in antibody-specific tasks,^6–8^ a multimodal AbLM could greatly expand our understanding of antibody-specific features and the broaden the potential applications of these models.

In summary, we present three key findings that will help inform future development and analysis of AbLMs. First, preferential masking of the non-templated CDR3 enhances antibody representation learning with both computational and data efficiency, achieving comparable or improved performance with 40% less training time. Second, VH/VL pairing is governed by learnable, non-random, sequence-inherent patterns, which could have implications for understanding antibody stability. Finally, the application of XAI highlights the fact that AbLMs are capable of learning meaningful biological patterns. Additionally, XAI enables interpretation of these complex patterns, offering potential insights into the fundamental immunological properties that underlie antibody development and function.

## METHODS

### Preferential Masking Implementation

The dynamic masking step of MLM is implemented in the HuggingFace data collator, where for each tokenized input sequence, a probability matrix is generated with the masking probability for each token index. The conventional probability matrix is a sequence-length vector of 0.15, which indicates uniform masking of the entire sequence at 15%. For preferential masking, we first construct a CDR mask for each sequence to reflect the region (FR or CDR1-3) in which the residue resides. Then, the masking probability *P*(*x*^*m*^) for each residue *x* was calculated using the following equation:

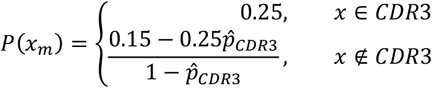

where 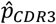refers to the proportion of residues in the sequence that reside in the CDR3.

For comparison with the conventional MLM strategy, an average masking probability of 15% across the entire sequence length was maintained. This is to ensure that comparisons are due to the modified masking strategy and not confounding factors such as increased overall masking. These probabilities were then constructed into a non-uniform probability matrix based on residue index. Construction of the CDR masks and subsequent calculation of preferential masking probabilities was done in a single initial pass over the entire dataset to eliminate redundant computation during model pre-training.

### Datasets

***Table 1*** gives a brief overview of the datasets used in this paper. Details on dataset construction can be found below.

**Table 1.** Dataset overview. Brief overview of the datasets used for model pre-training, testing, and downstream classification tasks.

| Dataset | Sources | Description of Use | Tables and Figures |
| --- | --- | --- | --- |
| A | Jaffe et al.<sup>25</sup><br>Hurtado et al.<sup>30</sup> | Base model pre-training (training and validation sets) | Fig. 2 |
| B | Jaffe et al.<sup>25</sup><br>Hurtado et al.<sup>30</sup> | Base model testing (per-residue inference) | Fig. 3, S1 |
| C | Jaffe et al.<sup>25</sup><br>Hurtado et al.<sup>30</sup> | Pair classification (same donors as pre-training, made by shuffling Dataset B) | Fig. 4 |
| D | Phad et al.<sup>31</sup><br>Data generated as part of this project | Pair classification (memory sequences from distinct donors from pre-training) | Fig. 4 |
| E | Raybould et al.<sup>32</sup><br>Phad et al.<sup>31</sup> | CoV-specificity classification and AttCAT analysis | Fig. 5, S2 |

#### Base Model Pre-training: Datasets A and B

For model pre-training, we used the largest publicly available dataset of natively paired antibody sequences, approximately 1.6 million pairs, from Jaffe et al.^25^ These sequences were isolated from circulating B-cells from healthy adult human donors (n=2), and were not enriched for any particular antigen-binding. We supplemented this original dataset with ∼400,000 paired B-cell sequences from the control dataset of Hurtado et al.,^30^ which were also isolated from circulating B-cells from healthy adult human donors (n=15). Raw sequences were annotated with abstar^27^ to extract the amino acid sequence of each V(D)J region. Label-encoded masks of the CDRs were constructed using the abutils package^27^ via semi-global alignment of V(D)J annotations to the full amino acid sequence. The data were filtered to remove duplicates and CDR mask alignment errors, and then clustered at 96% identity, resulting in 1,622,679 sequence pairs. The data was randomly split into training/validation/test sets at a ratio of 92:4:4, respectively, meaning that all donors are represented in all splits. The training and validation splits, which we refer to as Dataset A, were used for pre-training of the base models. The test split, which we refer to as Dataset B, was used for per-position inference.

#### Native Pair Classification: Datasets C and D

For the native pair classification task, healthy donor antibody sequences (not enriched for any particular antigen) were clustered at 95% identity, and non-native pairs were generated by sampling 50% of the sequences from each donor and randomly shuffling the heavy and light chains.^7^ Balanced classes for native vs. shuffled pairs allows for the splitting of training and test sets by donor. Due to light chain redundancy, a very small percentage (<0.1%) of the shuffled pairs were native; these were filtered out. The data was randomly split using a 5-fold Cross Validation (CV) with stratification.

Two native vs. shuffled pair datasets were created for this task. Dataset C contains naïve and memory B-cell sequences from the same donors as the pre-training data, and was created by shuffling Dataset B, resulting in 64,874 sequence pairs (32,437 of each class). Dataset D contains only memory B-cell sequences from different donors than Dataset A, and was created by shuffling paired memory B-cell sequences obtained from Phad et al.^31^ (n=2) and our own sequencing of PBMCs from healthy donors (n=3), resulting in 146,668 sequence pairs (73,334 of each class).

#### CoV-specificity Classification: Dataset E

For the specificity classification task, CoV antibody sequences were obtained from CoV-Ab-Dab.^32^ The negative set consisted of paired memory B-cell sequences from Phad et al.^31^ that were obtained pre-2020 to minimize the occurrence of false negative classifications. Amino acid sequences were clustered at 95% identity, resulting in 24,970 total sequence pairs (12,485 of each class), which we refer to as Dataset E. The data was randomly split using a 5-fold CV with stratification.

### Base Model Pre-training

We separately trained 2 models on Dataset A using an MLM objective: one using conventional uniformly random masking, and one using preferential masking. In both masking strategies, tokens were independently selected for prediction with an average probability of 15% across the entire input sequence length. From these, 80% are replaced with a mask token <mask>, 10% are replaced with a random token from the vocabulary, and the remaining 10% is left unchanged.^11^ Masking was performed dynamically^12^ to avoid encountering the same mask across epochs. For the preferential masking model, construction of the CDR masks and calculation of masking probabilities was done in a single initial pass over the entire dataset, stored, and reused in each epoch to eliminate redundant computation during model pre-training. Both models use an encoder-only ESM-2 architecture with 32 layers, 20 attention heads per layer, a hidden size of 960, and an intermediate size of 3840, resulting in both models having ∼350 million parameters.

The vocabulary contained 26 tokens: 1 for each of the proteinogenic amino acids, “X” for unnatural amino acids, and 5 special tokens: <pad>, <mask>, <unk>, <cls>, and <eos>. Inputs were heavy and light chain sequences concatenated by two <cls> tokens (***Fig. 1***) and padded to length 320 to accommodate the length of the longest sequence pair without truncation.

Both models were implemented using a slightly-modified HuggingFace transformers library^33^ and trained using DeepSpeed.^34^ Both models were trained using a total batch size of 256 for a total of 500,000 steps (∼85 epochs) on 8 NVIDIA A100 graphics processing units (**GPUs**), equating to ∼52 hours per model. The learning rate increased linearly to a peak of 1e-4 over the first 30,000 steps and decayed linearly thereafter.

The pre-training task of the preferential masking model was effectively harder than that of the uniform model, as increased CDR3 masking results in the prediction of more “difficult” residues with less context. Therefore, to compare training progression and monitor overfitting, evaluation loss for both models was calculated using cross-entropy loss under the uniform masking strategy. Overfitting was observed in both models (***Fig. 2A***), so the optimal checkpoints at 250,000 steps (∼43 epochs) and 350,000 steps (∼60 epochs) were chosen for downstream analyses.

### Classifier Model Training

A sequence classification head (single feedforward layer) was trained for binary classification of native vs. shuffled chain pairing (Datasets C and D), or CoV vs. healthy donor antibodies (Dataset E). Since we desire to evaluate the pre-training strategy via downstream applications of its output embeddings, the base model weights were frozen so as to not alter those embeddings during fine-tuning. Sequences were tokenized using the same tokenizer as pre-training, and no truncation was necessary as all sequences were shorter than the model’s maximum input length. Classifier models were trained for 50 epochs with a total batch size of 256, a peak learning rate of 1e-5, and a linear warm-up ratio of 0.1. For each classification task, 5 different shuffled, stratified splits of the data were generated using 5-fold CV, and a classifier head was trained independently on each of these splits for all base models. Training with a 5-fold CV was chosen to show variation based on the training/testing data.

Binary classifiers were evaluated using the following metrics: accuracy, area under to receiver operating characteristic curve (**AUC**), area under the precision-recall curve (**AUPR**), F1-score, and Matthews correlation coefficient (**MCC**). Test set accuracy and AUPR plots are averaged across 5 independent training runs with standard error. Metrics were calculated using scikit-learn and are based on Weights & Biases (wandb.ai) logging data.

### Specificity Classifier Model Interpretation

Attentive Class Activate Tokens (**AttCAT**) is a gradient-based attribution XAI method developed by Qiang et al.^23^ for visualizing which input tokens most inform a classification decision. Inspired by Gradient-weighted Class Activation Maps (**GradCAM**),^35^ AttCAT uses gradient information from all model layers and heads to quantify token impact. It is further adapted for the transformer architecture by integrating self-attention weights to capture global contextual information. The output is a set of “impact scores” that quantifies the informativeness of each input token for a specific target class. We chose this XAI method over other feature-attribution based methods or attention alone because AttCAT incorporates features, gradients, and self-attention into its score, giving a more comprehensive evaluation of all information flows through the model. Scores were normalized using standard scaling to facilitate comparison between models. Code was adapted from Song et al.^36^ to work with the ESM model architecture.

### Antibody Sequencing

De-identified peripheral blood mononuclear cells (**PBMC**) from healthy donors (n=3). PBMCs were first enriched for memory B-cells using the EasySep Human Memory B Cell Isolation Kit (Stemcell Technologies, catalog #17864) and sequenced using the 10x Genomics pipeline as described by Hurtado et al.^37^ Sequencing data were analyzed using Cell Ranger and custom Python scripts.

### Code and Data Availability

All code necessary to reproduce the results and figures in this paper is available through GitHub (github.com/brineylab/preferential-masking-paper). All data, including model weights and sequence datasets used for model training, validation, and testing are available through Zenodo (https://doi.org/10.5281/zenodo.13973760).

## ACKNOWLEDGEMENTS

The authors would like to thank Jonathan Hurtado and Simone Spandau for their help generating sequencing data (Dataset D) and Kenney Ng for helpful feedback on early manuscript drafts.

## AUTHOR CONTRIBUTIONS

Conceptualization: KN, BB

Model training and evaluation: KN

Manuscript preparation and revisions: KN, BB

## DECLARATION OF INTERESTS

BB is an equity shareholder in Infinimmune and a member of their Scientific Advisory Board.

**Figure S1.**
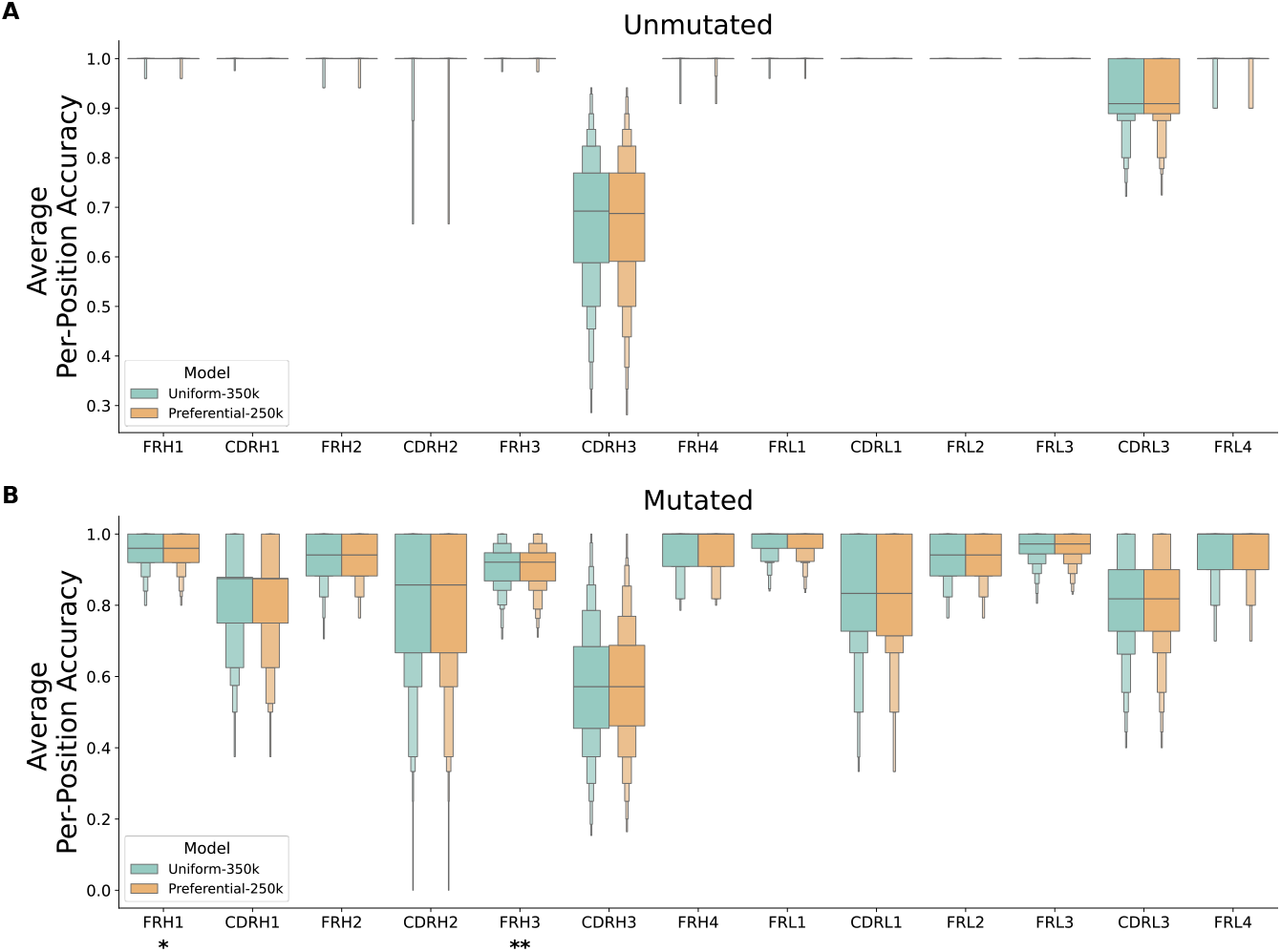
Per-position residue prediction accuracy. For 1000 unmutated (***A***) and mutated (***B***) test sequences, each residue was iteratively masked and predicted by both models (Uniform-350k and Preferential-250k). Mean prediction accuracy is plotted for each FR and CDR. Statistical significance for each region was calculated using a two-sided paired t-test with Bonferroni correction for multiple testing (14 regions).

**Figure S2.**
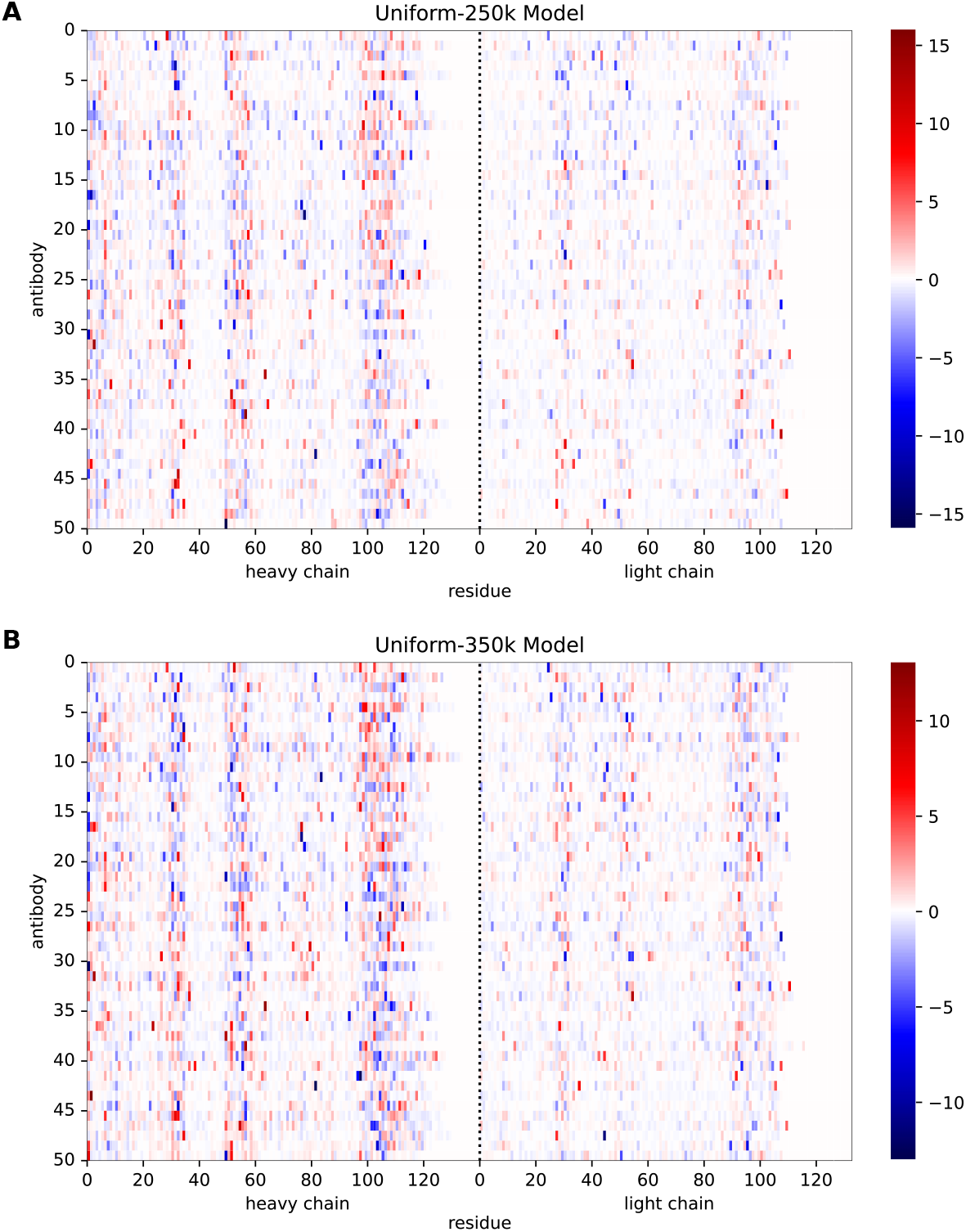
AttCAT analysis on the Uniform-250k and Uniform-350k CoV specificity classifier models. Normalized AttCAT impact scores with respect to the correct label class for the same 50 systematically chosen test sequences for the CoV specificity classifiers trained on the Uniform-250k (***A***) and Uniform-350k (***B***) base models. Sequences are sorted in ascending order by average prediction probability across the classifiers trained on all 3 base models.

